# Variants in regulatory elements of *PDE4D* associate with Major Mental Illness in the Finnish population

**DOI:** 10.1101/390518

**Authors:** Vishal Sinha, Liisa Ukkola-Vuoti, Alfredo Ortega-Alonso, Minna Torniainen-Holm, Sebastian Therman, Annamari Tuulio-Henriksson, Pekka Jylhä, Jaakko Kaprio, Iiris Hovatta, Erkki Isometsä, Tyrone D. Cannon, Jouko Lönnqvist, Tiina Paunio, Jaana Suvisaari, William Hennah

**Author notes:** Corresponding Author: William Hennah PhD, Institute for Molecular Medicine FIMM, P.O. Box 20, FI-00014, University of Helsinki, Helsinki, Finland.

## Abstract

We have previously reported a replicable association between variants at the *PDE4D* gene and familial schizophrenia in a Finnish cohort. In order to identify the potential functional mutations alluded to by these previous findings, we sequenced the 1.5Mb of the *PDE4D* genomic locus in 20 families (consisting of 96 individuals, and 79 independent chromosomes), followed by two stages of genotyping across 6,668 individuals from multiple Finnish cohorts for major mental illnesses. We identified 4,570 SNPs across the *PDE4D* gene, with 380 associated to schizophrenia (p≤0.05). Importantly, two of these variants, rs35278 and rs165940, are located at transcription factor binding sites, and displayed replicable association in the two-stage enlargement of the familial schizophrenia cohort, (combined statistics for rs35278 p=0.0012; OR=1.18, 95% CI 1.06-1.32; and rs165940 p=0.0016; OR=1.27, 95% CI 1.13-1.41). Further analysis using additional cohorts and endophenotypes revealed that rs165940 principally associates within the psychosis (p=0.025, OR=1.18, 95% CI 1.07-1.30) and cognitive domains of major mental illnesses (g-score p=0.044, beta=-0.033). Specifically, the cognitive domains represented verbal learning and memory (p=0.0091, beta=-0.044) and verbal working memory (p=0.0062, beta=-0.036). Moreover, expression data from the GTEx database demonstrated that rs165940 significantly correlates with the mRNA expression levels of *PDE4D* in the cerebellum (p-value=0.04; m-value=0.9), demonstrating a potential functional consequence for this variant. Thus, rs165940 represents the most likely functional variant for major mental illness at the *PDE4D* locus in the Finnish population, increasing risk broadly to psychotic disorders.

## 1. Introduction

The phosphodiesterase sub-family 4 are protein encoding genes belonging to the cyclic nucleotide phosphodiesterase (PDE) family, that play a key role in many important physiological processes through regulating and mediating of a number of cellular responses to extracellular signals(1). Members of the mammalian *PDE4* subfamily are evolutionary orthologues of the Drosophila learning and memory mutant Dunce(2). Flies carrying mutant Dunce display severe learning/memory phenotypes in different learning situations, showing reduced gene activity and deficits in olfactory learning and memory(3).

Although their involvement in human neuro-pathophysiology is currently unclear, several studies(3–18) have implicated *PDE4s* in psychiatric illnesses, particularly *PDE4D* and *PDE4B*. In a GWAS study of patient-related treatment response during antipsychotic therapy, three SNPs, in high linkage disequilibrium (LD), from *PDE4D* were found to significantly associate with mediating the effects of quetiapine(4). In acrodysostosis, a rare disorder characterized by intellectual disability, skeletal and neurological abnormalities, five different point mutations within the *PDE4D* gene have been identified as the genetic cause(5, 6). Furthermore, in a genome wide association study of neuroticism, a psychological trait reported to share genetic factors with both major depression and anxiety, found that one SNP in *PDE4D* associated with higher neuroticism(7). This observation has been replicated in two additional independent cohorts, but not in two other cohorts(7). Behavioural studies on *PDE4D* deficient mice reveal increased memory performance in the radial arm and water maze tasks, and object recognition tests(8). These results suggest that long-form PDE4D is important in mediation of memory and hippocampal neurogenesis through cAMP/CREB signaling, with reduced expression of PDE4D in the hippocampus enhancing memory. Genetic findings have also been noted with the *PDE4D* homolog *PDE4B*. With chromosomal translocations(9), SNP based haplotypes (10–12), and genome-wide SNP association (13) being observed in studies of schizophrenia. Moreover, genome-wide association studies (GWAS) have also identified PDE4B in the studies of anxiety(14).

Earlier studies have revealed that DISC1 interacts with the conserved regulatory UCR2 domain in both PDE4D and PDE4B, suggesting that DISC1 may modulate PDE4 catalytic activity(9). A recent study identified variants at the PDE4D locus as a potential risk modifier in the DISC1 translocation family, with follow-up analysis of two UK population-based cohorts (Generation Scotland and UK Biobank) finding suggestive association at PDE4D for affective disorders and related traits (17). In Finland, *PDE4D* and *PDE4B* were first identified as associated to schizophrenia in a study that deliberately set out to study known genes of DISC1 binding partners, as the DISC1 network had already been demonstrated to be of genetic importance within this large cohort for familial schizophrenia(19, 20). In this cohort of 476 families ascertained for schizophrenia, it was observed that haplotypes in both *PDE4D* (5q11.2-q12.1) and *PDE4B* (1p31.3) associated with schizophrenia in a replicable manner(18). For *PDE4D*, a haplotype comprised of the GGACA alleles of SNPs rs13190249, rs1120303, rs921942, rs10805515 and rs10514862, was observed to be significantly over-represented in affected individuals (p=0.00084). Moreover, the SNP rs1120303 also showed replicable association (p=0.021) in this dataset(18).

These associations already noted in the Finnish families were based on SNPs designed to tag the haplotypic structure of the genes of interest, and are thus not expected to be the functional mutations but instead only represent surrogate variants. Thus, here we sought to examine in detail *PDE4D*, which is the gene of the *PDE4* sub-family offering the more solid evidence for association to schizophrenia in the Finnish population, in order to identify any variants with potential functional consequences. This utilised a three-stage study design that would sequence the genomic locus of *PDE4D*, and two rounds of genotyping in ever increasing sample numbers from the familial schizophrenia cohort, so that variants of interest can be identified, verified, and replicated in an independent but identically ascertained cohort. This design was chosen as it is expected, based on the level of the prior observations (best p=0.00084, Tomppo et al. (18)), that any association with a single variant will not surpass the genome-wide significant threshold of 5.0 × 10^−8^, and thus replication of the findings is of paramount importance. Observations that passed through this three-stage design were then characterised further. We utilised seven other Finnish cohorts representing a range of major mental illness phenotypes, first individually and then jointly, to study the gene’s role in psychotic and mood disorders. Further, we studied neuropsychological endophenotype data that has been collected within the familial cohorts used here. See supplementary figure 1 for an illustration of the study work flow.

## 2. Materials and Methods

### 2.1. Samples

The samples studied came from multiple Finnish cohorts collected to study major psychiatric disorders using a number of different study designs, including familial, twin and population-based cohorts (Supplementary Table1), and phenotypes, including multiple diagnoses alongside neuropsychological endophenotypes. In total, this joint cohort consisted of 7,024 individuals for which 6,668 have been genotyped, including 1,909 psychiatrically healthy control samples. The individual cohorts have been described in great detail previously(21–26), and are briefly described, along with their abbreviations, in the supplementary materials. These samples include two familial cohorts, for schizophrenia (FSZ n= 2,818) and bipolar disorder (BPD n= 650), a sample of twin pairs concordant and discordant for schizophrenia (Twin n= 303), three population cohorts ascertained for different aspects of psychotic disorders (FEP n= 125; MMPN n= 449; HUPC n=383), a population cohort for anxiety disorders (Anx n= 823), and a sample of population controls (Controls n= 1,117).

In order to maximise the analytical potential of the cohorts studied here we combined all, except the 207 anxiety cases, into a joint analysis of two broad major mental illness related phenotypes. These dichotomous traits were derived from the diagnoses of each affected individual, where we could classify them as either having a psychotic disorder and/or mood disorder. The psychotic disorder phenotype included those individuals with any of the following diagnoses; schizophrenia, schizoaffective disorder, schizophrenia spectrum disorders, schizophreniform, psychotic disorders, and bipolar disorders and major depression only with a specification of including psychotic features. The mood disorder phenotype included individuals with the following phenotypes; bipolar disorder type I or type II, major depression, schizoaffective disorder, and psychotic disorders where major depressive or manic episodes had been present. In total, there were 1,896 people categorised with a psychotic disorder, and 1,227 with a mood disorder, 628 individuals could be categorised as cases under both phenotypes.

### 2.2. Sequencing, genotyping and SNP selection

The data to be investigated in this research have been generated through a three-stage replication study design to increase our power to detect true positives while lowering the amount of false positive findings, compared with using an overly conservative Bonferroni correction(27). Firstly, to identify variants in the familial schizophrenia cohort, we sequenced the 1.5Mb genomic region of the *PDE4D* gene (chr5:58254866-59793925; sequencing read depth=80.53; mean coverage=86.43) in a subset of 20 families using Illumia HiSeq2000, HiSeq1500 sequencing platform and Nimblegen SeqCap EZ 6Mb as the target enrichment kit. Sequencing data was analyzed using an in-house developed SAMtools-based bioinformatics pipeline (VCP) for quality control, short read alignment (using Burrows-Wheeler Aligner), duplicate removal, variant identification (with pileup) followed by variant annotation(28). The 20 families comprised 96 individuals (affected=42; Liability class (LC) LC1=28; LC2=12; LC3=2; LCs explained below), with 79 independent chromosomes for variant identification and segregation analysis. Identified variants were filtered by VCP according to the following: any variant call with a quality ratio of more than 0.8 was considered as a reference call and was filtered out. Calls with a quality ratio between 0.2 and 0.8 were considered to be heterozygous and calls below 0.2 to be homozygous variant calls. The filtered variants were aligned with bioinformatic predictions of function, with the major focus on their location in exonic and regulatory sites as predicted by UCSC genome browser build 19 (Supplementary Table 2). SNPs were selected for genotyping based on the following criteria. Firstly, SNPs displaying any evidence of LD|Linkage association in this small cohort (p≤0.05), and located in predicted functional areas: five exonic, one CpG, and eight TFBS. Additionally, SNPs were selected that showed Linkage p-value ≤0.05, and also had two other lines of evidence from TFBS, a score in the UCSC Brain Methylation track, or tentative association to an endophenotype for schizophrenia (social anhedonia) in our prior studies in a birth cohort from Northern Finland(29) (Supplementary Table 2). Of the 19 SNPs selected, seven dropped out due to inability to be fitted into a multiplex for Sequenom genotyping. Genotyping was performed using a Sequenom MassArray platform(30) according to manufacturer’s recommendations. Several quality control measures were applied across the genotyping stages, including high marker success rates (>95%), HWE p-value>0.01, optimal primer usage without leftover residues and no background signals in water. Stage one (S1) genotyping was performed in a sub-cohort of the schizophrenia families (n=1,122 individuals in 301 families) alongside population-based controls (n=323 individuals), in order to verify both the variant’s existence, if the variation was novel to this sequencing data, and independent association (p≤0.05) to schizophrenia. Those variants that were both verified and associated in this larger cohort were further taken for Stage two (S2) genotyping, which included the rest of the schizophrenia cohort (n=1,696), alongside the other major mental illness cohorts (Total n= 2,733) and additional population-based controls (n=794). The variant rs39672 displayed significant inconsistencies in its minor allele frequency between the controls of S1 and S2 (MAF S1=0.24; MAF S2=0.39: p=1.71×10^−11^; Supplementary Table 2) and was thus discarded, as there was no significant geographical or gender differences between these two stages that could reasonably account for such a difference.

### 2.3. Association analysis

Association analysis within the individual cohorts was carried out either using Pseudomarker(31) (family-based or twin cohorts) or PLINK(32) (population-based controls). Pseudomarker analyses test for single marker (two-point) association and linkage. Furthermore, Pseudomarker can utilize data of differing epidemiological design, helping us to combine familial studies with population controls, alongside singleton cases from other cohorts. It can handle missing data, even when the genotypes of the parents are unknown. The ‘association given linkage’ (LD|Linkage) option of the program was used to identify association for the SNPs with the diagnosis (p≤0.05), using the three main genetic models (dominant, recessive and additive) assuming incomplete penetrance. LD|Linkage corrects for the effect of any linkage within the families that may influence the observed association. The specific penetrance of alleles identified as associated in these cohorts was later calculated using the formula proposed by Wang et al (2006)(33). PLINK was used to analyse population-based cohorts, where the family structure was not known. The options --model and -- perm produced the p-values, using permutation tests to generate significance levels empirically.

In the family cohorts for schizophrenia and bipolar disorder different liability classes (LC) of the diagnoses were used in the analyses. The schizophrenia cohort liability classes (LCs) constitute of LC1 where individuals were diagnosed for schizophrenia only, LC2 added those individuals affected with schizoaffective disorder, LC3 added individuals with schizophrenia spectrum disorder, and LC4 added individuals with bipolar disorder or major depressive disorder. In the bipolar cohort, the liability classes (LC) are LC1 bipolar disorder type I, LC2 adds schizoaffective disorder, bipolar type, LC3 adds bipolar disorder type II, LC4 adds recurrent major depressive disorder. All other cohorts only used the single diagnostic criteria for which the cohort was ascertained (Supplementary Table 3-6). In the sequencing stage, association analysis was only performed using the broadest liability class possible, LC3, to maximise the number of cases (n=42) versus unaffected family members (n= 54).

### 2.4. Endophenotypes and QTDT

A total of 919 subjects from the schizophrenia (n = 811) and bipolar (n = 108) familial cohorts have been administered a comprehensive neuropsychological test battery, consisting of a series of well-validated, and internationally used, means to measure cognitive ability. From this test battery, a total of 14 quantitative neuropsychological variables with previous evidence of potential to use as an endophenotype were available(34–39). These endophenotypes that are included were: Verbal learning and memory derived from the Immediate, Short delay and Long delay recall tasks from the California Learning Test (CVLT) battery(40); Verbal skills from the Similarities and Vocabulary subtests of the Wechsler Adult Intelligence Scale–Revised (WAIS-R) battery(41); Visual working memory from the Visual span forward and backward, and Verbal working memory from the Digit span forward and backward tests from the Wechsler Memory Scale-Revised battery (WMS-R)(42); Information Processing from the Stroop colour task and colour-word task(43) and the Trail Making Test, parts A and B(44), and the Digit Symbol subtest from the Wechsler Adult Intelligence Scale– Revised (WAIS-R) battery(41). Most of the endophenotypes were normally distributed, however the Trail Making and Stroop tasks were transformed so that the scores represented speed (number of items/performance time). As the neuropsychological endophenotypes are related, they could be grouped into five first-order factors using factor analysis (Table2), and a second-order general ability factor (*g-factor*) based upon previous results(45) for cognition related traits. The factor analysis was performed with Mplus 7.3. We used confirmatory factor analysis with a maximum likelihood estimator and with robust standard errors using family as a cluster and affected versus not affected as a grouping variable. The fit of the model was acceptable (CFI= 0.932; RMSEA= 0.085).

The program QTDT(46) was used for the analysis of the endophenotypes, factors and *g-factor*. This method relies on variance components testing for any transmission distortion within the families. These 919 individuals contain 395 affected individuals, under the broadest diagnostic classifications for the two cohorts, and 524 unaffected relatives. Gender, age, and the broadest classification of affection (LC4) in both the schizophrenia and bipolar family cohorts were treated as covariates during the analysis. The orthogonal model (-ao) together with 100,000 permutations generated the empirical p-values. The orthogonal model allows for families of any size with or without parental genotypes, whereas permutation provides robust findings in the presence of stratification providing an empirical p-value for the observed difference in the trait. The strength for this analysis comes from the consistency of the findings with regard to related endophenotypes and factors. Due to the relatedness of these traits Bonferroni correction for multiple testing would be overly conservative, thus we report the uncorrected empirical p-values and highlight those that would survive multiple test correction for 14 endophenotypes (p=0.00357), five factors (p=0.010) and the single overall *g-factor* (p=0.05). Multiple regression using R software (with age, gender, affection status and family as covariates) was used to determine directionality of significant correlations between SNPs and endophenotypes through derivation of beta-estimates (β) (Supplementary Figure 2a, b).

### 2.6. Linkage Disequilibrium (LD) analysis

In order to investigate the relatedness between our identified variants, and between our observations and those of the previous study(18), the identified variants from this study were mapped onto the previously determined LD haplotypes of *PDE4D* using the Haploview program(47). For the latter, the D′ haplotype structure of the SNPs previously studied was manually enforced onto this data with Hedrick’s Multiallelic D′ (48) determined between the haplotype and the two identified SNPs. However, for the correlation between our two identified SNPs the r^2^ LD was calculated.

## 3. Results

The sequencing of *PDE4D* in 20 families identified 4,570 variants, of which 380 variants associated with a broad schizophrenia diagnosis (p≤0.05). Using multiple levels of information 12 variants were genotyped in the first stage (S1), of which four were both verified and remained significant in their association to schizophrenia in this enlarged cohort, and therefore taken forward for replication in the second genotyping stage (S2). Two potential regulatory SNPs were identified to significantly replicate in their association to schizophrenia (Table 1, Figure 1, Supplementary Table 4), and were thus studied within the combined familial schizophrenia cohort; rs35278 (LC3 additive; p=0.0012; OR=1.18, ±95% CI 1.06-1.32) and rs165940 (LC3 additive; p=0.0016; OR=1.27, ±95% CI 1.13-1.41). These two SNPs are in relatively high linkage disequilibrium, r^2^ =0.66. No gender or geographical-based differences were observed in our analyses.

**Figure 1:**
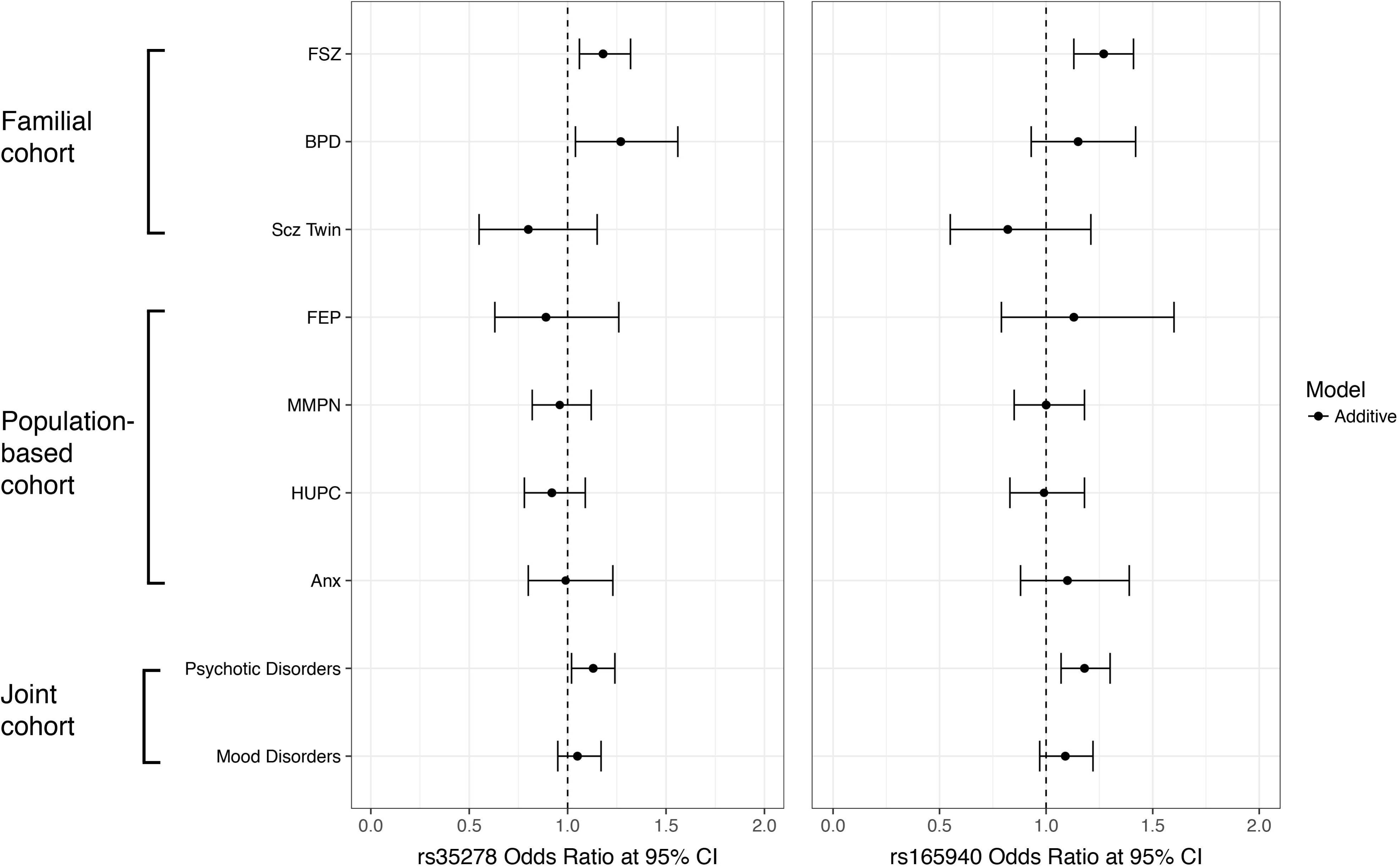
Odds Ratios, and their respective 95% confidence intervals, for the two SNPs across the cohorts studied and the joint analysis. All plots represent the observations for the additive genetic model. For the schizophrenia family cohort only the finding for liability class 3 is shown, while for the familial bipolar disorder cohort only liability class 2 is shown.

**Table 1:** Replicated observations of association across the three-stage study design for two *PDE4D* SNPs within the Finnish familial schizophrenia cohort. P-values, odds ratios and confidence intervals are shown for liability class 3 under an additive genetic model. Values for other models and classes are provided in Supplementary Table 3.

| SNP | Allele | Sequencing<br>P-value | S1<br>P-value | S2<br>P-value | Combined<br>(S1+S2)<br>P-value | Odds Ratio<br>(95% CI) | MAF SCZ<br>Families | MAF<br>Control |
| --- | --- | --- | --- | --- | --- | --- | --- | --- |
| rs35278 | T>G | **0.011** | **0.033** | **0.0074** | **0.0012** | **1.18**<br>**(1.06 - 1.32)** | 0.446 | 0.384 |
| rs165940 | A>T | **0.047** | **0.0082** | **0.0084** | **0.0016** | **1.27**<br>**(1.13 - 1.41)** | 0.360 | 0.289 |

Association analysis in the other major mental illness cohorts revealed that rs35278 significantly associates within the familial bipolar disorder cohort with broadening liability classes, but most significantly with LC2, bipolar type I and schizoaffective, bipolar type, (LC2 additive p=0.032; OR=1.27, ±95% CI 1.04-1.56) (Supplementary Table 5). We observe that both of our variants are partially penetrant in these ascertained familial cohorts, with rs35278 displaying low (FSZ=0.15, BPD=0.21), and rs165940 (FSZ=0.36, BPD=0.50) displaying moderate penetrance in these families for the respective LCs providing the best evidence of association. Neither SNP associated within the other individual cohorts (Supplementary Table 6), however, the joint cohort analysis highlights the involvement of both SNPs with a broad diagnosis of any psychotic disorder (rs35278 additive p=0.023, OR=1.13, ±95% CI 1.02-1.24; rs165940 additive p=0.025, OR=1.18, ±95% CI 1.07-1.30) (Supplementary Table 6).

Follow-up analysis of these two SNPs using quantitative neuropsychological endophenotypes demonstrated that both SNPs provide evidence for association to overall measure representing a g-score (rs35278 p=0.043, β=-0.025; rs165940 p=0.044, β=-0.033) (Table 2). When investigating the sub-domains of cognition through use of factors only rs165940 continued to display association levels that would survive Bonferroni correction for the number of factors tested. The minor allele, T, significantly associates with decreased performance in the factors representing verbal working memory (p=0.0062, β=-0.036), and verbal learning and memory (p=0.0091, β=-0.044)(Table 2). Analysis of the individual endophenotypes shows that under the factor verbal learning and memory, the T allele associates with decreased scores on the immediate recall task (p=0.0032, β=-0.65) (Table2).

**Table 2:** Empirical p-values from the association analysis of the two *PDE4D* SNPs and the quantitative neuropsychological endophenotypes, their factors, and an overall general score for cognition.

| Endophenotype | rs35278 | rs165940 | Factor | rs35278 | rs165940 | g-factor |  |
| --- | --- | --- | --- | --- | --- | --- | --- |
|  |  |  |  |  |  | rs35278 | rs165940 |
| WAIS-R; Vocabulary | 0.056 | 0.17 | Verbal skills | 0.015 | 0.063 | 0.043 | 0.044 |
| WAIS-R; Similarities | 0.024 | 0.15 |  |  |  |  |  |
| WAIS-R; Digit symbol | 0.043 | 0.10 | Information Processing | 0.10 | 0.15 |  |  |
| Stroop color naming, time | 0.41 | 0.56 |  |  |  |  |  |
| Stroop color-word task, time | 0.80 | 0.48 |  |  |  |  |  |
| Trail Making A, time | 0.18 | 0.17 |  |  |  |  |  |
| Trail Making B, time | 0.24 | 0.31 |  |  |  |  |  |
| WMS-R; Digit span forward | 0.33 | 0.048 | Verbal working memory | 0.011 | 0.0062 |  |  |
| WMS-R; Digit span backward | 0.0062 | 0.0092 |  |  |  |  |  |
| WMS-R; Visual span forward | 0.60 | 0.16 | Visual working memory | 0.25 | 0.18 |  |  |
| WMS-R; Visual span backward | 0.60 | 0.98 |  |  |  |  |  |
| CVLT Immediate recall | 0.013 | 0.0032 | Verbal learning and memory | 0.013 | 0.0091 |  |  |
| CVLT; Short delay recall | 0.010 | 0.014 |  |  |  |  |  |
| CVLT; Long delay recall | 0.067 | 0.044 |  |  |  |  |  |

P-values $\leq 0.05$ for variables and factors are marked in bold.
P-values that surpass the Bonferroni multiple test correction for either the number of endophenotypes (14), the number of factors (5) or g-factor (1) tested are in bold and italics.

Since both the SNPs have a potential regulatory function based on their locations being predicted at transcription factor binding sites, we checked the extent of their functional consequences using tissue specific expression quantitative trait loci (eQTL) from the GTEx portal(49). Only rs165940 showed significant association with expression changes in *PDE4D* within the brain, specifically the cerebellum region (p-value=0.04, m-value=0.9) (Supplementary Figure 3). Significant modulation of expression of *PDE4D* by rs165940 is also detected in the oesophagus, heart and prostate tissues, the meta-analysis over all tissues gives a p-value of 0.00000017.

## Discussion

Through the use of a three-stage sequencing and genotyping approach to identify, validate, and replicate potential functional mutations at the *PDE4D* gene, we have identified two SNPs of principal interest to the aetiology of major mental illness in Finland. Both SNPs displayed replicable association in the familial schizophrenia cohort, are located in predicted transcription factor binding sites, and are in relatively high LD with each other (r^2^=0.66). Significant association with these SNPs was observed across liability classes and genetic models, the relevant phenotype was further refined using additional cohorts (see below) while the fact that all genetic models displayed associations suggest that the additive model is the most likely. The issue of LD makes it difficult to discern which of the two SNPs is the most likely functional mutation. The higher allele frequency of rs35278 gives it a greater power for detection in the association analyses, thus, use of additional cohorts for major mental illness highlighted only rs35278 as also associating with bipolar disorder. However, when the cohorts were combined to study psychotic and mood disorders as a whole, both SNPs again displayed significant association, to psychotic disorders. The minor alleles of both SNPs (rs35278 G allele, rs165940 T allele) were significantly enriched in the disorders. The use of neuropsychological endophenotypes indicates that both SNPs display significant association to an overall measure of cognition, while analysis of factors and individual endophenotypes shows both SNPs displaying association, but with only T allele of rs165940 being significantly so, to reduced scores in the factors representing verbal learning and memory, and verbal working memory and the endophenotype immediate recall. Through the analysis of the functional consequences of these SNPs on the gene expression levels of *PDE4D* within the GTEx database we gained extra insight that could help to specifically separate these two variants, with only rs165940 significantly associating with gene expression levels in *PDE4D*, making it the most likely functional mutation of the two SNPs. These expression level differences could be identified in the brain, the cerebellum, but also other tissues, such as oesophagus, heart and prostate. While these expression changes indicate the potential functional consequence of rs165940 on PDE4D, these are not necessarily in tissues of direct relevance to the phenotypes being studied. Although there is growing evidence for the involvement of the cerebellum in major mental illness(50–53), our identification of association to learning and memory would more traditionally be associated with functions of the hippocampus. It can be noted that the eQTL status of rs165940 on PDE4D in hippocampus is approaching significance but the GTEx data size is smaller compared to that of the cerebellum (hippocampus n=111, cerebellum n=154; Supplementary Figure 3). As sample sizes increase, this may become truly significant.

Since this study is an extension of one that had previously implicated *PDE4D* in this familial cohort for schizophrenia, we determined whether our current findings are in concordance with our prior observations(18). Thus we mapped our SNPs onto the SNPs and haplotypes of *PDE4D* previously used to study the common haplotypic background of the gene, as determined by D’ LD. The SNPs are located in approximately the same physical area as the haplotype, being within the same intron, intron 3 of the longest isoform of PDE4D. Furthermore, both SNPs are within some degree of D′ LD (D’=0.43,0.47) with the haplotype (Supplementary Figure 4a, b), which, although modest, suggests some relationship between these observations and the original observation.

It is important to note that the approach taken in this study design is both a strength, as it enables us to directly replicate our findings using an identically ascertained cohort, but also a weakness. The initial sequencing step only used 96 related individuals, consisting of 79 independent chromosomes, meaning that the full scope of variation at the *PDE4D* locus within these families has not been determined, with rarer mutations (MAF<0.01) in these families likely to have been missed. However, our prior evidence at *PDE4D* implied that any mutation would be common (frequency > 5%), as the associating haplotype allele had a frequency in affected individuals from this cohort of 0.40 (control frequency=0.28), and the SNP which showed some association had a minor allele frequency (MAF) of 0.13 (control MAF=0.19). The high frequency of rs165940, not just in the cohorts studied here, Finnish familial schizophrenia cases (MAF=0.34, 78% of FSZ families carry one or more copies of the risk T allele), and psychiatrically healthy controls (MAF=0.29), but also in population genomic cohorts(54) for the Finnish population (MAF=0.32) and the non-Finnish Europeans (MAF=0.28), does in turn suggest that these variants could have been detected by large scale genome-wide approaches, and yet, to date, this is not the case. While current genetic evidence identified in familial cohorts for schizophrenia is markedly different to those identified through population-based study designs, it is worth noting that the latest consortia-based genome-wide association study has identified *PDE4B* in its study of schizophrenia(13). This is part of the same sub-family as *PDE4D* and was also identified as associating to schizophrenia in the Finnish familial schizophrenia cohort used here. The high frequency of the *PDE4D* SNPs, combined with their low odds ratio/effect size, would imply that with further increases in sample size *PDE4D* could also be identified through population-based approaches. Whereas our three-stage study design has allowed us to identify significant variants through replication, rather than being dependent on reaching the genome wide significance threshold of 5.0 × 10^−8^.

Our findings strongly support the role of *PDE4D* in psychiatric disorders, with replicable association in familial schizophrenia in Finland. Further characterisation suggested that it plays a role in both psychosis and cognitive endophenotypes of major mental illnesses. In particular we demonstrate that the SNP rs165940, through its association pattern and being identified as an eQTL for the *PDE4D* gene, make it the principal variant of interest for being the functional mutation at this locus. The variant rs165940 is located in a transcription factor binding site positioned in an intron of the longer isoforms of PDE4D but upstream of two short isoforms. From the data used here it cannot be discerned if there are specific PDE4D transcripts being altered by this locus, yet given its location it would be expected that it any eQTL effect this SNP has on PDE4D may well be isoform specific. Thus, further studies into the functional consequences of this variant are essential.

## Supporting information

## Acknowledgements

NGS library preparation, enrichment, sequencing and sequence analysis were performed by the Institute for Molecular Medicine Finland FIMM Technology Centre, University of Helsinki. We sincerely thank FIMM’s sequencing and genotyping unit, especially Pekka Ellonen and Kati Donner for their efforts in producing the data. We also thank Sarang Talwelkar and Disha Malani for their input in improving the figures. This study has been funded by the Academy of Finland (128504, 259589 and 265097), MC-ITN EU-FP7 (607616) and Finnish Cultural Foundation (Ingrid, Toini and Olavi Martelius Grant 2018) for WH, Sigrid Juselius Foundation for JL, and Jalmari and Rauha Ahokkas Foundation for VS. The funders had no further role in the study design; nor in the collection, analysis, and interpretation of data, in the writing of the report, nor in the decision to submit the paper for publication.

## Contributors

VS and WH wrote the manuscript text and prepared the manuscript tables and figures; WH designed the study; TP, JS, JL, IH, PJ, EI, ATH, ST, TDC and JK provided access to samples and data; VS, LUV, WH, MTH, ST and AOA performed the analysis. All authors have reviewed the manuscript and approved the final version to be published.

## Conflict of Interest

WH has a co-appointment at Orion Pharmaceuticals. All other authors declare no conflicts of interest.

## Notes

#### Summary of Updates

Manuscript updated

## References

1. Lugnier C. Cyclic nucleotide phosphodiesterase (PDE) superfamily: a new target for the development of specific therapeutic agents. Pharmacology & therapeutics. 2006;109(3):366–98.

2. Davis RL. Physiology and biochemistry of Drosophila learning mutants. Physiological reviews. 1996;76(2):299–317.

3. Davis RL, Cherry J, Dauwalder B, Han PL, Skoulakis E. The cyclic AMP system and Drosophila learning. Molecular and cellular biochemistry. 1995;149-150:271–8.

4. Clark SL, Souza RP, Adkins DE, Aberg K, Bukszar J, McClay JL, et al. Genome-wide association study of patient-rated and clinician-rated global impression of severity during antipsychotic treatment. Pharmacogenetics and genomics. 2013;23(2):69–77.

5. Lindstrand A, Grigelioniene G, Nilsson D, Pettersson M, Hofmeister W, Anderlid BM, et al. Different mutations in PDE4D associated with developmental disorders with mirror phenotypes. Journal of medical genetics. 2014;51(1):45–54.

6. Lee H, Graham JM, Jr., Rimoin DL, Lachman RS, Krejci P, Tompson SW, et al. Exome sequencing identifies PDE4D mutations in acrodysostosis. American journal of human genetics. 2012;90(4):746–51.

7. Shifman S, Bhomra A, Smiley S, Wray NR, James MR, Martin NG, et al. A whole genome association study of neuroticism using DNA pooling. Molecular psychiatry. 2008;13(3):302–12.

8. Li YF, Cheng YF, Huang Y, Conti M, Wilson SP, O’Donnell JM, et al. Phosphodiesterase-4D knock-out and RNA interference-mediated knock-down enhance memory and increase hippocampal neurogenesis via increased cAMP signaling. The Journal of neuroscience : the official journal of the Society for Neuroscience. 2011;31(1):172–83.

9. Millar JK, Pickard BS, Mackie S, James R, Christie S, Buchanan SR, et al. DISC1 and PDE4B are interacting genetic factors in schizophrenia that regulate cAMP signaling. Science (New York, NY). 2005;310(5751):1187–91.

10. Fatemi SH, King DP, Reutiman TJ, Folsom TD, Laurence JA, Lee S, et al. PDE4B polymorphisms and decreased PDE4B expression are associated with schizophrenia. Schizophrenia research. 2008;101(1-3):36–49.

11. Numata S, Ueno S, Iga J, Song H, Nakataki M, Tayoshi S, et al. Positive association of the PDE4B (phosphodiesterase 4B) gene with schizophrenia in the Japanese population. Journal of psychiatric research. 2008;43(1):7–12.

12. Pickard BS, Thomson PA, Christoforou A, Evans KL, Morris SW, Porteous DJ, et al. The PDE4B gene confers sex-specific protection against schizophrenia. Psychiatric genetics. 2007;17(3):129–33.

13. Pardinas AF, Holmans P, Pocklington AJ, Escott-Price V, Ripke S, Carrera N, et al. Common schizophrenia alleles are enriched in mutation-intolerant genes and in regions under strong background selection. Nature genetics. 2018;50(3):381–9.

14. Meier S, Trontti K, Als TD, Laine M, Pedersen MG, Bybjerg-Grauholm J, et al. Genome-wide Association Study of Anxiety and Stress-related Disorders in the iPSYCH Cohort. bioRxiv. 2018.

15. Siuciak JA, McCarthy SA, Chapin DS, Martin AN. Behavioral and neurochemical characterization of mice deficient in the phosphodiesterase-4B (PDE4B) enzyme. Psychopharmacology. 2008;197(1):115–26.

16. Kuroiwa M, Snyder GL, Shuto T, Fukuda A, Yanagawa Y, Benavides DR, et al. Phosphodiesterase 4 inhibition enhances the dopamine D1 receptor/PKA/DARPP-32 signaling cascade in frontal cortex. Psychopharmacology. 2012;219(4):1065–79.

17. Ryan NM, Lihm J, Kramer M, McCarthy S, Morris SW, Arnau-Soler A, et al. DNA sequence-level analyses reveal potential phenotypic modifiers in a large family with psychiatric disorders. Molecular psychiatry. 2018.

18. Tomppo L, Hennah W, Lahermo P, Loukola A, Tuulio-Henriksson A, Suvisaari J, et al. Association between genes of Disrupted in schizophrenia 1 (DISC1) interactors and schizophrenia supports the role of the DISC1 pathway in the etiology of major mental illnesses. Biological psychiatry. 2009;65(12):1055–62.

19. Hennah W, Varilo T, Kestila M, Paunio T, Arajarvi R, Haukka J, et al. Haplotype transmission analysis provides evidence of association for DISC1 to schizophrenia and suggests sex-dependent effects. Human molecular genetics. 2003;12(23):3151–9.

20. Hennah W, Tomppo L, Hiekkalinna T, Palo OM, Kilpinen H, Ekelund J, et al. Families with the risk allele of DISC1 reveal a link between schizophrenia and another component of the same molecular pathway, NDE1. Human molecular genetics. 2007;16(5):453–62.

21. Ekholm JM, Pekkarinen P, Pajukanta P, Kieseppa T, Partonen T, Paunio T, et al. Bipolar disorder susceptibility region on Xq24-q27.1 in Finnish families. Molecular psychiatry. 2002;7(5):453–9.

22. Kaprio J, Koskenvuo M, Rose RJ. Population-based twin registries: illustrative applications in genetic epidemiology and behavioral genetics from the Finnish Twin Cohort Study. Acta geneticae medicae et gemellologiae. 1990;39(4):427–39.

23. Cannon TD, Kaprio J, Lonnqvist J, Huttunen M, Koskenvuo M. The genetic epidemiology of schizophrenia in a Finnish twin cohort. A population-based modeling study. Archives of general psychiatry. 1998;55(1):67–74.

24. Mantere O, Saarela M, Kieseppa T, Raij T, Mantyla T, Lindgren M, et al. Anti-neuronal anti-bodies in patients with early psychosis. Schizophrenia research. 2018;192:404–7.

25. Aaltonen K, Naatanen P, Heikkinen M, Koivisto M, Baryshnikov I, Karpov B, et al. Differences and similarities of risk factors for suicidal ideation and attempts among patients with depressive or bipolar disorders. Journal of affective disorders. 2016;193:318–30.

26. Donner J, Pirkola S, Silander K, Kananen L, Terwilliger JD, Lonnqvist J, et al. An association analysis of murine anxiety genes in humans implicates novel candidate genes for anxiety disorders. Biological psychiatry. 2008;64(8):672–80.

27. van den Oord EJ, Sullivan PF. A framework for controlling false discovery rates and minimizing the amount of genotyping in the search for disease mutations. Human heredity. 2003;56(4):188–99.

28. Sulonen AM, Ellonen P, Almusa H, Lepisto M, Eldfors S, Hannula S, et al. Comparison of solution-based exome capture methods for next generation sequencing. Genome biology. 2011;12(9):R94.

29. Ortega-Alonso A, Ekelund J, Sarin AP, Miettunen J, Veijola J, Jarvelin MR, et al. Genome-Wide Association Study of Psychosis Proneness in the Finnish Population. Schizophrenia bulletin. 2017;43(6):1304–14.

30. Jurinke C, van den Boom D, Cantor CR, Koster H. Automated genotyping using the DNA MassArray technology. Methods in molecular biology (Clifton, NJ). 2001;170:103–16.

31. Gertz EM, Hiekkalinna T, Digabel SL, Audet C, Terwilliger JD, Schaffer AA. PSEUDOMARKER 2.0: efficient computation of likelihoods using NOMAD. BMC bioinformatics. 2014;15:47.

32. Purcell S, Neale B, Todd-Brown K, Thomas L, Ferreira MA, Bender D, et al. PLINK: a tool set for whole-genome association and population-based linkage analyses. American journal of human genetics. 2007;81(3):559–75.

33. Wang Y, Ottman R, Rabinowitz D. A method for estimating penetrance from families sampled for linkage analysis. Biometrics. 2006;62(4):1081–8.

34. Cannon TD, Hennah W, van Erp TG, Thompson PM, Lonnqvist J, Huttunen M, et al. Association of DISC1/TRAX haplotypes with schizophrenia, reduced prefrontal gray matter, and impaired short- and long-term memory. Archives of general psychiatry. 2005;62(11):1205–13.

35. Hennah W, Tuulio-Henriksson A, Paunio T, Ekelund J, Varilo T, Partonen T, et al. A haplotype within the DISC1 gene is associated with visual memory functions in families with a high density of schizophrenia. Molecular psychiatry. 2005;10(12):1097–103.

36. Paunio T, Tuulio-Henriksson A, Hiekkalinna T, Perola M, Varilo T, Partonen T, et al. Search for cognitive trait components of schizophrenia reveals a locus for verbal learning and memory on 4q and for visual working memory on 2q. Human molecular genetics. 2004;13(16):1693–702.

37. Tuulio-Henriksson A, Arajarvi R, Partonen T, Haukka J, Varilo T, Schreck M, et al. Familial loading associates with impairment in visual span among healthy siblings of schizophrenia patients. Biological psychiatry. 2003;54(6):623–8.

38. Tuulio-Henriksson A, Haukka J, Partonen T, Varilo T, Paunio T, Ekelund J, et al. Heritability and number of quantitative trait loci of neurocognitive functions in families with schizophrenia. American journal of medical genetics. 2002;114(5):483–90.

39. Wedenoja J, Loukola A, Tuulio-Henriksson A, Paunio T, Ekelund J, Silander K, et al. Replication of linkage on chromosome 7q22 and association of the regional Reelin gene with working memory in schizophrenia families. Molecular psychiatry. 2008;13(7):673–84.

40. Delis D KJ, Kaplan E, Ober B. California verbal learning test (CVLT). San Antonio: The Psychological Corporation. 1987.

41. D. W. Manual for the Wechsler adult intelligence scale-revised (WAIS-R). San Antonio, TX: The Psychological Corporation. 1981.

42. D. W. WMS-R: Wechsler memory scale-revised. San Antonio, TX: Psychological Corporation. 1987.

43. Golden CJ FS. Stroop color and word test. 1978.

44. RM R. Trail Making Test: Manual for administration and scoring. : Reitan Neuropsychology Laboratory. 1986.

45. Liisa Ukkola-Vuoti MT-H, Alfredo Ortega-Alonso, Vishal Sinha, Annamari Tuulio-Henricksson, Tiina Paunio, Jouko Lönnqvist, Jaana Suvisaari, William Hennah. Gene expression changes related to immune processes associate with cognitive endophenotypes of schizophrenia. *Under review Neuropsychopharmacology & Biological Psychiatry. 2018.

46. Abecasis GR, Cardon LR, Cookson WO. A general test of association for quantitative traits in nuclear families. American journal of human genetics. 2000;66(1):279–92.

47. Barrett JC, Fry B, Maller J, Daly MJ. Haploview: analysis and visualization of LD and haplotype maps. Bioinformatics (Oxford, England). 2005;21(2):263–5.

48. Hedrick PW. Gametic disequilibrium measures: proceed with caution. Genetics. 1987;117(2):331–41.

49. The Genotype-Tissue Expression (GTEx) project. Nature genetics. 2013;45(6):580–5.

50. He H, Luo C, Luo Y, Duan M, Yi Q, Biswal BB, et al. Reduction in gray matter of cerebellum in schizophrenia and its influence on static and dynamic connectivity. Human brain mapping. 2018.

51. Peters H, Shao J, Scherr M, Schwerthoffer D, Zimmer C, Forstl H, et al. More Consistently Altered Connectivity Patterns for Cerebellum and Medial Temporal Lobes than for Amygdala and Striatum in Schizophrenia. Frontiers in human neuroscience. 2016;10:55.

52. Kim DJ, Kent JS, Bolbecker AR, Sporns O, Cheng H, Newman SD, et al. Disrupted modular architecture of cerebellum in schizophrenia: a graph theoretic analysis. Schizophrenia bulletin. 2014;40(6):1216–26.

53. O’Halloran CJ, Kinsella GJ, Storey E. The cerebellum and neuropsychological functioning: a critical review. Journal of clinical and experimental neuropsychology. 2012;34(1):35–56.

54. Zerbino DR, Achuthan P, Akanni W, Amode M R, Barrell D, Bhai J, et al. Ensembl 2018. Nucleic Acids Research. 2018;46(D1):D754–D61.

