## Supplementary material for "Variants in regulatory elements of *PDE4D* associate with Major Mental Illness in the Finnish population"

^*^Corresponding Author:

William Hennah PhD

**Supplementary Materials and Methods**

**Descriptions of the Cohorts Used in this Study**

***Finnish Familial Schizophrenia cohort (FSZ)***

The familial cohort for schizophrenia comprises 670 families, that contain 2,818 individuals with available DNA, including 1,309 individuals classified as affected according to increasingly inclusive liability classes (LC), based on the Diagnostic and Statistical Manual of Mental Disorders, fourth edition (DSM-IV)(1). LC1 constitutes schizophrenia only (affected=911), LC2 added those individuals affected with schizoaffective disorder (+154), LC3 added individuals with schizophrenia spectrum disorder (+160), and LC4 added individuals with bipolar disorder or major depressive disorder (+84). We sequenced 20 families consisting of 96 individuals (affected=42) and 79 independent chromosomes for variation discovery within this ascertained cohort. For genotyping, we randomly split the family samples into two non-overlapping identically ascertained sub-samples to be analysed as verification and replication data sets. The verification set consists of 1,122 individuals from 301 families (stage 1 (S1)), and the replication set contains 1,553 individuals from 369 families (stage 2 (S2)). The combined sample set contains 2,818 individuals from 670 families, this includes both stages (S1+S2) and an additional 143 individuals who were genotyped in stage 2 but belonged to the families analysed in stage 1. These families were recruited both from an internal isolate with an increased incidence of schizophrenia(2), as well as from the rest of Finland(3). There was no difference in the geographical origins between the two stages used here.

***Bipolar disorder cohort (BPD)***

This Bipolar disorder cohort(4) consists of 181 families, with 650 individuals, ascertained for bipolar disorder, but allows for the study of a broad spectrum of related diagnoses as determined based on the DSM-IV. These were studied based on the same liability class approach as described for the familial schizophrenia cohort, where the liability classes are (LC1) bipolar disorder type I (affected=206); (LC2) adds schizoaffective disorder, bipolar type (+29); (LC3) adds bipolar disorder type II (+14); (LC4) adds recurrent major depressive disorder (+20).

Both the familial schizophrenia and bipolar disorder make up the “THL’s psychiatric family collections” biobank. Additional information about this collection is available through the websites of the National Institute for Health and Welfare(5, 6).

***Twin cohort (Scz Twin)***

The twin cohort subsample used here consists of 195 twin pairs that are principally discordant for schizophrenia(7, 8). Briefly, the data set was collected by identifying discordant psychosis-related register data among like-sexed pairs originally identified from the Finnish National Population Register, and diagnostically interviewing them. This cohort consists of 303 individuals, of whom 77 have been diagnosed for schizophrenia, 187 individuals are known controls, and the diagnosis is unknown for 39 individuals. From these twin pairs 80 (32 case pairs (one or both twins affected), and 48 control pairs (neither affected)) are monozygotic, 109 (40 case pairs, and 69 control pairs) are dizygotic, and 6 pairs with only one twin available (4 cases, 2 controls). The programs for association analysis could not fully accommodate twin pair status into the analysis, thus only one member of each monozygotic pair has been used in analysis, selection of which individual per pair primarily chose the case member of the pair, or if both were cases, then choosing randomly. Dizygotic pairs were treated as full siblings.

***First Episode Psychosis cohort (FEP)***

The First Episode Psychosis (9) cohort are patients with first-episode psychosis, aged 18-40 years, recruited from the catchment area of the Helsinki University Hospital and the city of Helsinki, Finland, between November 2010 and September 2015. All primary psychotic disorders were included, whereas substance-induced psychoses and psychotic disorders due to a general medical condition were excluded. The inclusion criterion was a score of at least 4 in the items assessing delusions (Unusual Thought Content) or hallucinations in the Brief Psychiatric Rating Scale - Extended (BPRS-E)(10). The Research version of the Structured Clinical Interview for DSM-IV was used for diagnostic assessment. Medical records from all psychiatric treatment contacts were also reviewed for the final diagnostic assessment according to DSM-IV diagnostic criteria. This cohort contains 125 individuals, with 81 diagnosed for psychosis, 39 controls and 5 with an unknown diagnostic criteria.

***MMPN cohort (MMPN)***

The MMPN cohort comprises three separate but comparable clinical cohorts collected from psychiatric in- and out-patients from the metropolitan area of southern Finland: Jorvi Bipolar study (JoBS), The Vantaa depression study (VDS), and the Vantaa Primary Care Depression Study (PC-VDS) (11). In all studies, patient sampling was based on screening for BD (JoBS), MDD (VDS), and depressive disorders (PC-VDS). The cohorts were collected from districts that provide secondary care psychiatric services for psychiatric patients (JoBS) or primary care (PC-VDS). To diagnose mood and other axis I disorders and to assess psychotic symptoms, the Research version of the Structured Clinical Interview for DSM-IV was used for diagnostic assessment. In total, the MMPN cohort comprised 429 affected individuals, of which 33.1% are males and 66.9% are females.

***Helsinki University Psychiatry Consortium cohort (HUPC)***

HUPC is a population-based cohort consisting of 383 individuals with diagnoses from a broad range of disorders, including severe depressive disorder or bipolar disorder with suicidal tendencies, alcohol use disorder, borderline personality disorder traits, and childhood physical abuse. These individuals are from the Helsinki metropolitan area, including the municipalities of Helsinki, Espoo, Vantaa, Kauniainen, Kerava, and Kirkkonummi. The study diagnoses were based on the patients’ clinical diagnoses assigned by the attending physicians, the validity of which was critically evaluated as described in Aaltonen et al. (12). The diagnostic classification followed the International Statistical Classification of Diseases and Related health Problems, 10^th^ Revision, Diagnostic Criteria for Research. Each patient was given hierarchically a diagnosis regarded as the most severe and pervasive over the lifetime, in which severe depressive, bipolar affective, and psychotic disorders were given precedence over other conditions. Bipolar affective disorder was classified into type I and II disorders, based on the established Finnish practice of applying Diagnostic and Statistical Manual of Mental Disorders (DSM) – compatible classification in national treatment guidelines. Information on psychotic illness was based on the clinical main diagnosis and on cases with mood disorder, on information on diagnosis of psychotic mania or psychotic depression.

***Anxiety cohort (Anx)***

This anxiety disorder cohort(13) is a population-based cohort derived from the much larger Health 2000 cohort from all over Finland that is designed to study many different aspects of health in the general Finnish population. The cohort used here consists of 207 cases, who were either diagnosed, according to DSM-IV(1), or sub-threshold, according to Composite International Diagnostic Interview (CIDI)(14), for any anxiety disorder. This disorder classification includes panic disorder, generalised anxiety disorder, social phobia, agoraphobia, and phobia not specified otherwise. Furthermore, this cohort contains an additional 566 controls selected based on the absence of any neuropsychiatric diagnoses. These controls have been matched according to gender, age (±1 year), and hospital catchment area.

***Controls***

The control samples were derived from the same Health 2000 (H2000) population-based cross-sectional health study as the anxiety cohort. From the 8,028 total participants in this cohort, we randomly selected 1,117 psychiatrically healthy individuals, and were not included as controls in the anxiety cohort. This cohort consists of individuals from all over Finland, with 51.7% males and 48.3% females.

**Additional References:**

1.	DSM-IV-TR: American Psychiatric Association; 2000. 982 p.

2.	Hovatta I, Varilo T, Suvisaari J, Terwilliger JD, Ollikainen V, Arajarvi R, et al. A genomewide screen for schizophrenia genes in an isolated Finnish subpopulation, suggesting multiple susceptibility loci. American journal of human genetics. 1999;65(4):1114-24.

3.	Ekelund J, Hovatta I, Parker A, Paunio T, Varilo T, Martin R, et al. Chromosome 1 loci in Finnish schizophrenia families. Human molecular genetics. 2001;10(15):1611-7.

4.	Ekholm JM, Pekkarinen P, Pajukanta P, Kieseppa T, Partonen T, Paunio T, et al. Bipolar disorder susceptibility region on Xq24-q27.1 in Finnish families. Molecular psychiatry. 2002;7(5):453-9.

5.	<https://thl.fi/en/web/thl-biobank/for-researchers/sample-collections/health-2000-and-2011-surveys>.

6.	<https://thl.fi/en/web/thl-biobank/for-researchers/sample-collections/thl-psychiatric-family-collections-1994-2008>.

7.	Kaprio J, Koskenvuo M, Rose RJ. Population-based twin registries: illustrative applications in genetic epidemiology and behavioral genetics from the Finnish Twin Cohort Study. Acta geneticae medicae et gemellologiae. 1990;39(4):427-39.

8.	Cannon TD, Kaprio J, Lonnqvist J, Huttunen M, Koskenvuo M. The genetic epidemiology of schizophrenia in a Finnish twin cohort. A population-based modeling study. Archives of general psychiatry. 1998;55(1):67-74.

9.	Mantere O, Saarela M, Kieseppa T, Raij T, Mantyla T, Lindgren M, et al. Anti-neuronal anti-bodies in patients with early psychosis. Schizophrenia research. 2018;192:404-7.

10.	Ventura J LD, Nuechterlein K, Liberman R, Green M, Shaner A. Brief psychiatric rating scale (BPRS), expanded version (4.0): Scales, anchor points, and administration manual. Int J Methods Psychiatr Res. 1993;3:227-43.

11.	Soronen P, Mantere O, Melartin T, Suominen K, Vuorilehto M, Rytsala H, et al. P2RX7 gene is associated consistently with mood disorders and predicts clinical outcome in three clinical cohorts. American journal of medical genetics Part B, Neuropsychiatric genetics : the official publication of the International Society of Psychiatric Genetics. 2011;156b(4):435-47.

12.	Aaltonen K, Naatanen P, Heikkinen M, Koivisto M, Baryshnikov I, Karpov B, et al. Differences and similarities of risk factors for suicidal ideation and attempts among patients with depressive or bipolar disorders. Journal of affective disorders. 2016;193:318-30.

13.	Donner J, Pirkola S, Silander K, Kananen L, Terwilliger JD, Lonnqvist J, et al. An association analysis of murine anxiety genes in humans implicates novel candidate genes for anxiety disorders. Biological psychiatry. 2008;64(8):672-80.

14.	Robins LN, Wing J, Wittchen H, et al. The composite international diagnostic interview: An epidemiologic instrument suitable for use in conjunction with different diagnostic systems and in different cultures. Archives of general psychiatry. 1988;45(12):1069-77.

15.	Ortega-Alonso A, Ekelund J, Sarin AP, Miettunen J, Veijola J, Jarvelin MR, et al. Genome-Wide Association Study of Psychosis Proneness in the Finnish Population. Schizophrenia bulletin. 2017;43(6):1304-14.

16.	The Genotype-Tissue Expression (GTEx) project. Nature genetics. 2013;45(6):580-5.

17.	Tomppo L, Hennah W, Lahermo P, Loukola A, Tuulio-Henriksson A, Suvisaari J, et al. Association between genes of Disrupted in schizophrenia 1 (DISC1) interactors and schizophrenia supports the role of the DISC1 pathway in the etiology of major mental illnesses. Biological psychiatry. 2009;65(12):1055-62.

18.	Barrett JC, Fry B, Maller J, Daly MJ. Haploview: analysis and visualization of LD and haplotype maps. Bioinformatics (Oxford, England). 2005;21(2):263-5.

**Supplementary Table 1:** Number of individuals from each cohort for which DNA was available for this study

| **Cohort** | **Sample Size** | **Gender**  **(M/F)** | **Cases*** | **Controls** | **Endo**  **phenotypes** | **Psychotic**  **Disorders**** | **Mood Disorders**** |
| --- | --- | --- | --- | --- | --- | --- | --- |
| FSZ | 2818 | 1417/1401 | 1309 | 0 | 811 | 1298 | 238 |
| Controls | 1117 | 578/539 | 0 | 1117 | 0 | 0 | 0 |
| Anx | 823 | 297/526 | 207 | 566 | 0 | 0 | 0 |
| BPD | 650 | 308/342 | 269 | 0 | 108 | 199 | 267 |
| MMPN | 449 | 152/297 | 449 | 0 | 0 | 116 | 449 |
| HUPC | 383 | 132/251 | 383 | 0 | 0 | 126 | 248 |
| Scz Twin | 303 | 137/166 | 77 | 187 | 0 | 78 | 8 |
| FEP | 125 | 78/47 | 81 | 39 | 0 | 79 | 17 |
| Total | 6,668 | 3,099/3,569 | 2,775 | 1,909 | 919 | 1,896 | 1,227 |

*Number of cases according to the broadest diagnostic category used for that cohort, see supplementary materials and methods for more details.

**Case individuals classified into these joint categories are not mutually exclusive.

**Supplementary Table 2:** Findings from the three stages of analysis of the familial schizophrenia cohort. The table shows the variants identified through sequencing 96 individuals, their location, and frequency. Additional columns display the bioinformatic predictions of function for that location as annotated through the table function of UCSC, and the HWE p-values, linkage and association results obtained from Pseudomarker across all stages and the final combined analysis.

Table is separate file; Supplementary_Table_2.xlsx

* Minor allele frequencies (MAF) are reported using three different categories, annotated in three different columns. Primarily the allele frequency of variants has been determined only in the founders of this sub-sample, so as to avoid inflation of the MAF through including related individuals. These are annotated in the MAF (Founders) column. However, as not all parents were genotyped, some variants were identified exclusive of the founders. The MAF of these variants has been calculated in the whole cohort and annotated in the column MAF (Whole Cohort). The frequencies reported in this column are expected to be artificially higher due to being calculated in related individuals. Variants with no reported MAF were monomorphic for this cohort, but for the non-reference allele. These 240 variants are labelled in a column headed “Monomorphic

** Social Anhedonia GWAS p-values from Ortega-Alonso et al (15). Other psychosis proneness traits were analysed in that publication. However, social anhedonia is more established as a psychosis proneness measure, and was thus the only one used here.

**Supplementary Table 3:** Minor allele frequency (MAF) and Hardy-Weinberg equilibrium p-value (HWE) of our identified SNPs across all the cohorts. These summary statistics have been derived using --freq and --hwe options from the PLINK software, and --nonfounders as an additional option where applicable.

| **SNP** | **Info** | **Schizophrenia** | **H2000** | **Anxiety** | **Bipolar Disorder** | **MMPN** | **HUPC** | **Twins** | **FEP** |
| --- | --- | --- | --- | --- | --- | --- | --- | --- | --- |
| rs35278 | MAF | 0.45 | 0.38 | 0.39 | 0.42 | 0.38 | 0.37 | 0.36 | 0.38 |
|  | HWE | 0.55 | 1.00 | 0.46 | 0.81 | 0.26 | 0.89 | 0.79 | 0.47 |
| rs165940 | MAF | 0.36 | 0.29 | 0.31 | 0.31 | 0.32 | 0.30 | 0.26 | 0.34 |
|  | HWE | 0.75 | 0.58 | 0.62 | 0.52 | 0.73 | 0.80 | 0.22 | 0.39 |

**Supplementary Table 4:** Full association results from the three-stage study design within the Finnish familial schizophrenia cohort. Showing results for Pseudomarker LD given Linkage across the four Liability classes (LC) and three genetic models tested.

| **Stage** | **SNP** | **Info** | **LC1rec** | **LC1add** | **LC1dom** | **LC2rec** | **LC2add** | **LC2dom** | **LC3rec** | **LC3add** | **LC3dom** | **LC4rec** | **LC4add** | **LC4dom** |
| --- | --- | --- | --- | --- | --- | --- | --- | --- | --- | --- | --- | --- | --- | --- |
| S1 | rs35278 | p-val | 0.24 | 0.094 | 0.088 | **0.029** | **0.036** | **0.043** | **0.048** | **0.033** | **0.046** | 0.12 | **0.037** | **0.045** |
|  |  | OR | 1.59 | 1.28 | 1.13 | 1.72 | 1.32 | 1.13 | 1.81 | 1.36 | 1.17 | 1.72 | 1.33 | 1.15 |
|  |  | 95% CI | 0.94 - 2.69 | 1.10 - 1.64 | 0.80 - 1.60 | 1.05 - 2.84 | 1.04 - 1.67 | 0.81 - 1.57 | 1.13 - 2.95 | 1.08 - 1.71 | 0.85 - 1.61 | 1.07 - 2.77 | 1.06 - 1.71 | 0.85 - 1.57 |
|  | rs165940 | p-val | 0.34 | 0.061 | **0.043** | **0.038** | **0.021** | **0.016** | 0.057 | **0.0082** | **0.010** | 0.27 | **0.023** | **0.021** |
|  |  | OR | 1.58 | 1.22 | 1.20 | 1.84 | 1.30 | 1.27 | 1.92 | 1.40 | 1.34 | 1.82 | 1.31 | 1.28 |
|  |  | 95% CI | 0.87 - 2.92 | 0.95 - 1.57 | 0.86 - 1.68 | 1.06 - 3.28 | 1.02 - 1.65 | 0.93 - 1.74 | 1.13 - 3.37 | 1.11 - 1.77 | 0.99 - 1.82 | 1.07 - 3.17 | 1.04 - 1.64 | 0.95 - 1.73 |
| S2 | rs35278 | p-val | 0.26 | **0.0072** | **0.0044** | 0.29 | 0.090 | 0.071 | **0.0082** | **0.0074** | **0.0070** | **0.0079** | **0.022** | **0.018** |
|  |  | OR | 1.22 | 1.21 | 1.34 | 1.17 | 1.16 | 1.26 | 1.22 | 1.18 | 1.27 | 1.26 | 1.21 | 1.30 |
|  |  | 95% CI | 0.94 - 1.58 | 1.05- 1.39 | 1.08 - 1.66 | 0.91 - 1.50 | 1.02 - 1.33 | 1.03 - 1.54 | 0.97 - 1.54 | 1.04 - 1.34 | 1.06 - 1.54 | 1.01 - 1.58 | 1.07 - 1.37 | 1.08 - 1.57 |
|  | rs165940 | p-val | **0.046** | **0.0084** | **0.0066** | 0.11 | 0.059 | **0.044** | **0.027** | **0.035** | **0.032** | **0.026** | **0.021** | **0.017** |
|  |  | OR | 1.34 | 1.12 | 1.37 | 1.34 | 1.23 | 1.27 | 1.40 | 1.25 | 1.29 | 1.41 | 1.27 | 1.32 |
|  |  | 95% CI | 0.97 - 1.83 | 0.97 - 1.29 | 1.12 - 1.67 | 0.99 - 1.81 | 1.07 - 1.41 | 1.06 - 1.53 | 1.05 - 1.85 | 1.09 - 1.42 | 1.08 - 1.53 | 1.07 - 1.86 | 1.11 - 1.44 | 1.11 - 1.57 |
| S1+S2 | rs35278 | p-val | 0.13 | **0.0035** | **0.0026** | **0.030** | **0.0097** | **0.010** | **0.0010** | **0.0012** | **0.0014** | **0.0022** | **0.0011** | **0.0012** |
|  |  | OR | 1.27 | 1.08 | 1.23 | 1.30 | 1.17 | 1.18 | 1.33 | 1.18 | 1.20 | 1.33 | 1.19 | 1.21 |
|  |  | 95% CI | 1.03 - 1.58 | 1.05 - 1.32 | 1.03 - 1.46 | 1.06 - 1.59 | 1.05 - 1.30 | 1.00 - 1.39 | 1.09 - 1.61 | 1.06 - 1.32 | 1.03 - 1.40 | 1.10 - 1.61 | 1.07 - 1.32 | 1.04 - 1.41 |
|  | rs165940 | p-val | 0.060 | **0.0016** | **0.0010** | **0.017** | **0.0028** | **0.0016** | **0.0038** | **0.0016** | **0.0012** | **0.012** | **0.0034** | **0.0013** |
|  |  | OR | 1.32 | 1.24 | 1.31 | 1.24 | 1.24 | 1.27 | 1.46 | 1.27 | 1.30 | 1.41 | 1.26 | 1.30 |
|  |  | 95% CI | 1.01 - 1.72 | 1.10 - 1.40 | 1.11 - 1.54 | 0.96 - 1.59 | 1.11 - 1.40 | 1.09 - 1.49 | 1.15 - 1.86 | 1.13 - 1.41 | 1.12 - 1.50 | 1.12 - 1.79 | 1.13 - 1.41 | 1.12 - 1.50 |

p-values <0.05 are marked in bold, the cut-off we used to look for replication across stages.

**Supplementary Table 5:** Full association results from the Finnish familial bipolar disorder cohort. Showing results for Pseudomarker LD given Linkage across the four Liability classes (LC) and three genetic models tested.

| **Cohort** | **SNP** | **Info** | **LC1rec** | **LC1add** | **LC1dom** | **LC2rec** | **LC2add** | **LC2dom** | **LC3rec** | **LC3add** | **LC3dom** | **LC4rec** | **LC4add** | **LC4dom** |
| --- | --- | --- | --- | --- | --- | --- | --- | --- | --- | --- | --- | --- | --- | --- |
| BPD | rs35278 | p-val | 0.92 | 0.051 | **0.032** | 0.41 | **0.032** | **0.015** | 0.25 | **0.036** | **0.017** | 0.77 | **0.048** | **0.025** |
|  |  | OR | 1.52 | 1.29 | 1.32 | 1.48 | 1.27 | 1.31 | 1.41 | 1.25 | 1.29 | 1.37 | 1.26 | 1.35 |
|  |  | 95% CI | 1.03 - 2.19 | 1.04 - 1.60 | 0.96 - 1.84 | 1.03 - 2.09 | 1.04 - 1.56 | 0.97 - 1.79 | 0.99 - 1.99 | 1.02 - 1.51 | 0.96 - 1.74 | 0.97 - 1.91 | 1.04 - 1.52 | 1.01 - 1.81 |
|  | rs165940 | p-val | 0.97 | 0.19 | 0.13 | 0.42 | 0.24 | 0.16 | 0.25 | 0.31 | 0.2 | 0.32 | 0.38 | 0.23 |
|  |  | OR | 1.17 | 1.20 | 1.30 | 1.06 | 1.15 | 1.24 | 1.00 | 1.11 | 1.19 | 1.00 | 1.10 | 1.17 |
|  |  | 95% CI | 0.68 - 1.90 | 0.96 - 1.50 | 0.96 - 1.75 | 0.63 - 1.71 | 0.93 - 1.42 | 0.93 - 1.65 | 0.60 - 1.61 | 0.90 - 1.37 | 0.90 - 1.58 | 0.61 - 1.59 | 0.90 - 1.34 | 0.89 - 1.53 |

p-values <0.05 are marked in bold.

**Supplementary Table 6:** Association findings across genetic models from the cohorts with a single diagnostic category, and the joint disorder analysis.

| **Cohort** | **Diagnosis** | **SNP** | **Info** | **Recessive** | **Additive** | **Dominant** |
| --- | --- | --- | --- | --- | --- | --- |
| Scz Twin | Schizophrenia | rs35278 | p-val | 0.34 | 0.24 | 0.26 |
|  |  |  | OR | 0.77 | 0.80 | 0.74 |
|  |  |  | 95% CI | 0.33 - 1.58 | 0.55 - 1.15 | 0.45 - 1.24 |
|  |  | rs165940 | p-val | 0.43 | 0.30 | 0.31 |
|  |  |  | OR | 1.06 | 0.82 | 0.72 |
|  |  |  | 95% CI | 0.40 - 2.35 | 0.55 - 1.21 | 0.43 - 1.18 |
| FEP | Psychotic Disorders | rs35278 | p-val | 0.62 | 1.00 | 0.48 |
|  |  |  | OR | 0.89 | 0.89 | 0.85 |
|  |  |  | 95% CI | 0.41 - 1.72 | 0.63 - 1.26 | 0.53 - 1.40 |
|  |  | rs165940 | p-val | 0.30 | 0.85 | 0.66 |
|  |  |  | OR | 1.42 | 1.13 | 1.07 |
|  |  |  | 95% CI | 0.64 - 2.85 | 0.79 - 1.60 | 0.67 - 1.72 |
| MMPN | Psychotic Disorders | rs35278 | p-val | 0.41 | 0.43 | 1.00 |
|  |  |  | OR | 0.84 | 0.96 | 1.00 |
|  |  |  | 95% CI | 0.61 - 1.14 | 0.82 - 1.12 | 0.80 - 1.25 |
|  |  | rs165940 | p-val | 1.00 | 1.00 | 0.85 |
|  |  |  | OR | 1.01 | 1.00 | 1.00 |
|  |  |  | 95% CI | 0.69 - 1.47 | 0.85 - 1.18 | 0.81 - 1.24 |
| HUPC | Psychotic Disorders | rs35278 | p-val | 0.66 | 0.72 | 0.64 |
|  |  |  | OR | 0.86 | 0.92 | 0.93 |
|  |  |  | 95% CI | 0.61 - 1.19 | 0.78 - 1.09 | 0.73 - 1.17 |
|  |  | rs165940 | p-val | 0.85 | 1.00 | 0.66 |
|  |  |  | OR | 0.90 | 0.99 | 1.05 |
|  |  |  | 95% CI | 0.58 - 1.36 | 0.83 - 1.18 | 0.84 - 1.32 |
| Anx | Anxiety Disorders (Inc. sub-threshold) | rs35278 | p-val | 0.85 | 0.77 | 0.66 |
|  |  |  | OR | 1.09 | 0.99 | 0.94 |
|  |  |  | 95% CI | 0.71 - 1.63 | 0.80 - 1.23 | 0.69 - 1.30 |
|  |  | rs165940 | p-val | 0.35 | 0.77 | 0.52 |
|  |  |  | OR | 1.21 | 1.10 | 1.10 |
|  |  |  | 95% CI | 0.71 - 1.98 | 0.88 - 1.39 | 0.81 - 1.50 |
| Psychotic Disorders | All Psychotic Disorders | rs35278 | p-val | **0.0037** | **0.023** | 0.057 |
|  |  |  | OR | 1.22 | 1.13 | 1.14 |
|  |  |  | 95% CI | 1.02 - 1.45 | 1.02 - 1.24 | 0.99 - 1.31 |
|  |  | rs165940 | p-val | **0.028** | **0.025** | **0.032** |
|  |  |  | OR | 1.29 | 1.18 | 1.21 |
|  |  |  | 95% CI | 1.03 - 1.60 | 1.07 - 1.30 | 1.06 - 1.37 |
| Mood Disorders | All Mood Disorders | rs35278 | p-val | 0.28 | 0.81 | 0.81 |
|  |  |  | OR | 1.08 | 1.05 | 1.07 |
|  |  |  | 95% CI | 0.88 - 1.32 | 0.95 - 1.17 | 0.92 - 1.25 |
|  |  | rs165940 | p-val | 0.054 | 1.00 | 0.79 |
|  |  |  | OR | 1.17 | 1.09 | 1.09 |
|  |  |  | 95% CI | 0.91 - 1.50 | 0.97 - 1.22 | 0.94 - 1.26 |

p-values <0.05 are marked in bold.

**Supplementary Figure 1:** Schematic overview of the PDE4D analysis workflow.

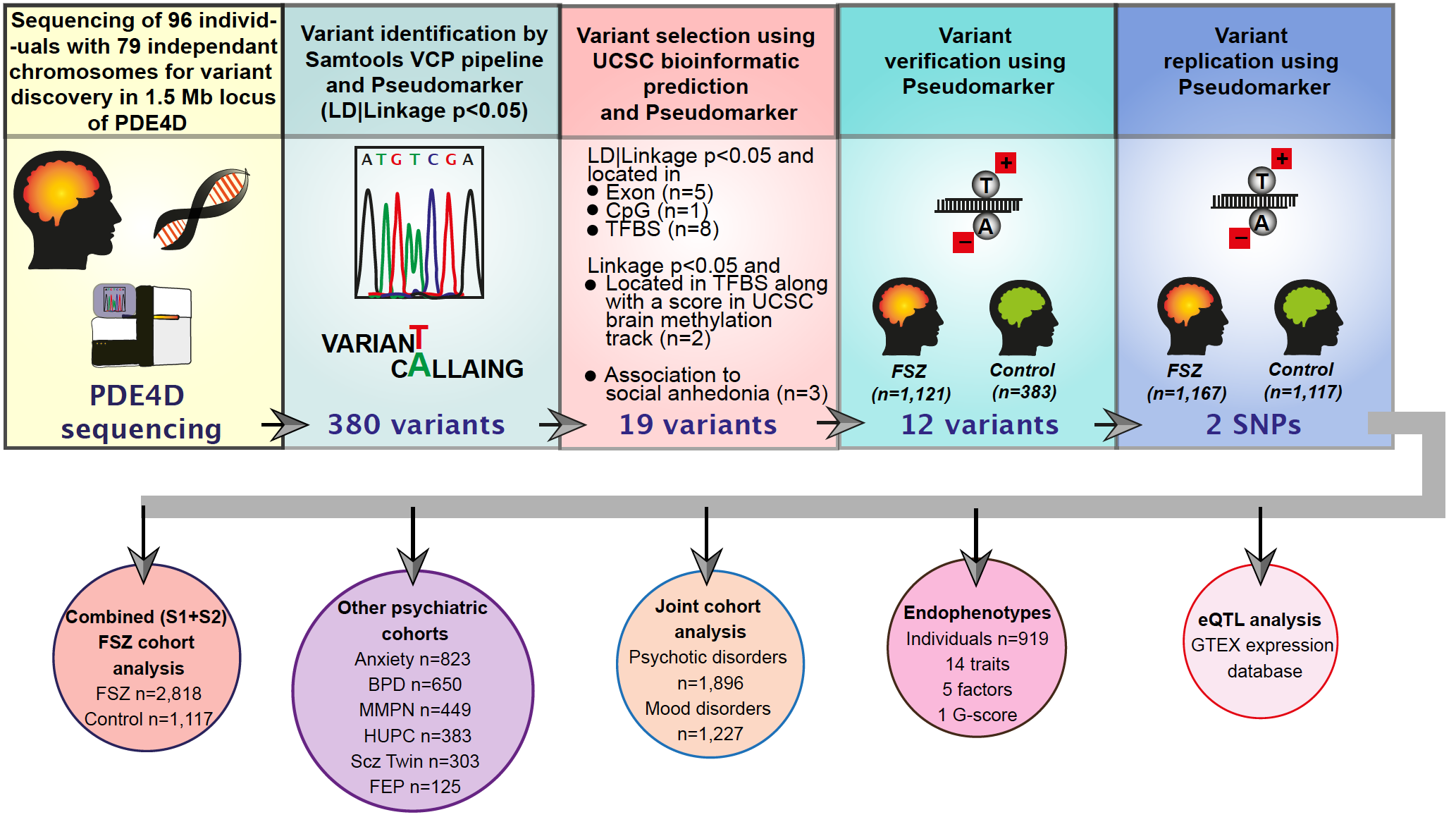

**Supplementary Figure 2:** The association analyses for endophenotypes were performed using the QTDT program. Age, gender and affection status were used as covariates. Association p-values (QTDT) and beta estimates (multiple regression) of significant correlations has been provided for SNPs (a) rs165940, (b) rs35278 and endophenotypes.

**a)**

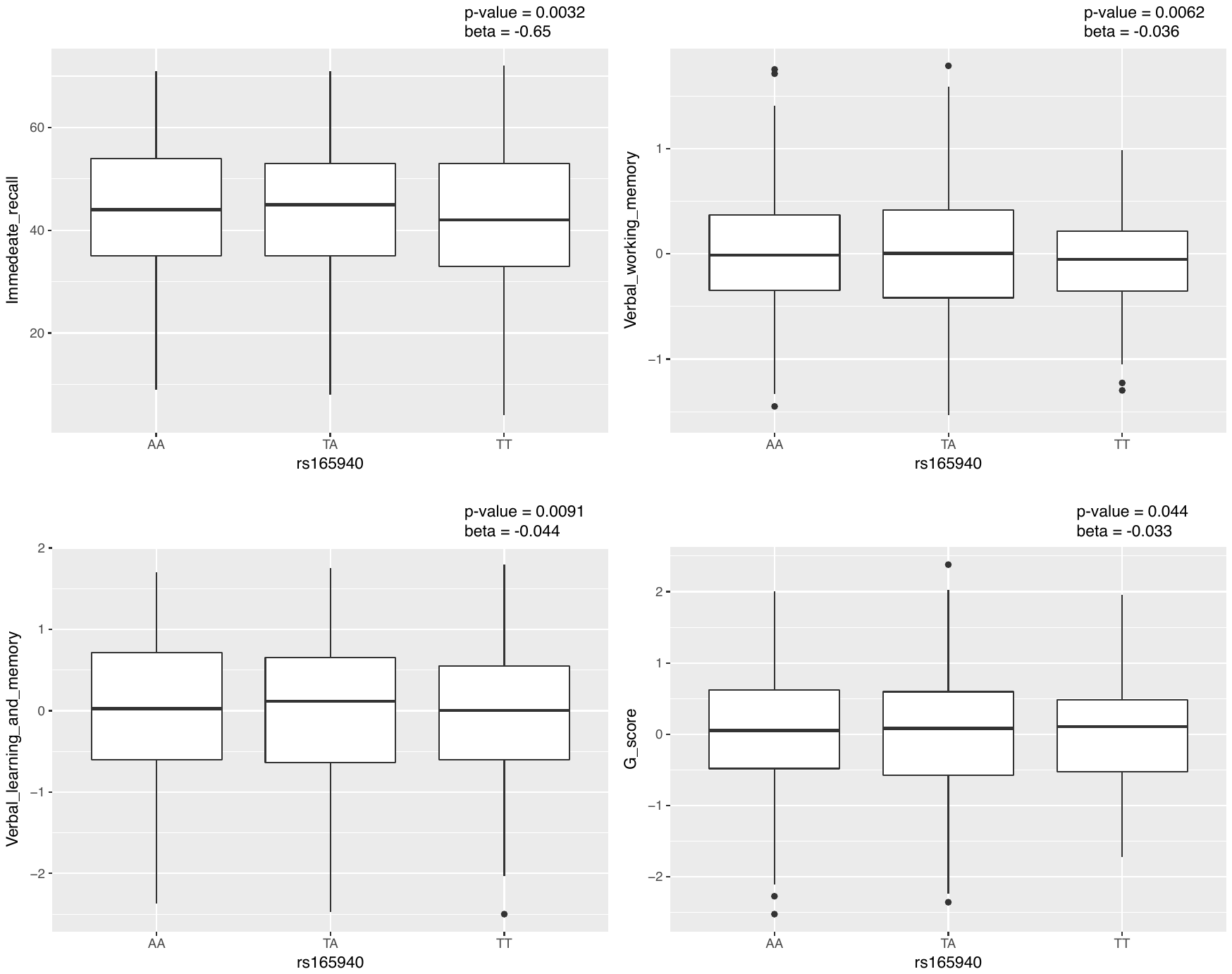

**b)**

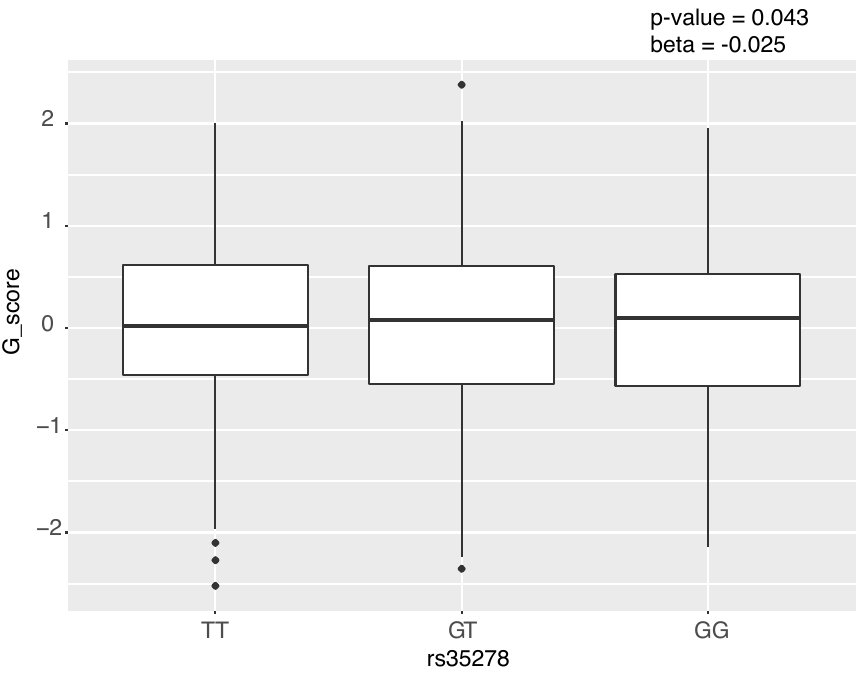

**Supplementary Figure 3:** eQTL analysis of rs165940. Figure from the GTEx database (16)(figure used under general permission, accessed on 20.11.2018) showing the gene expression changes in *PDE4D* associated with rs165940 across tissue types. Red Arrow: rs165940 alters the expression of *PDE4D* in the cerebellum (p-value=0.04; m-value=0.9). Red Box: rs165940 also alters PDE4D expression in oesophagus - mucosa, heart - atrial appendage, and prostate.

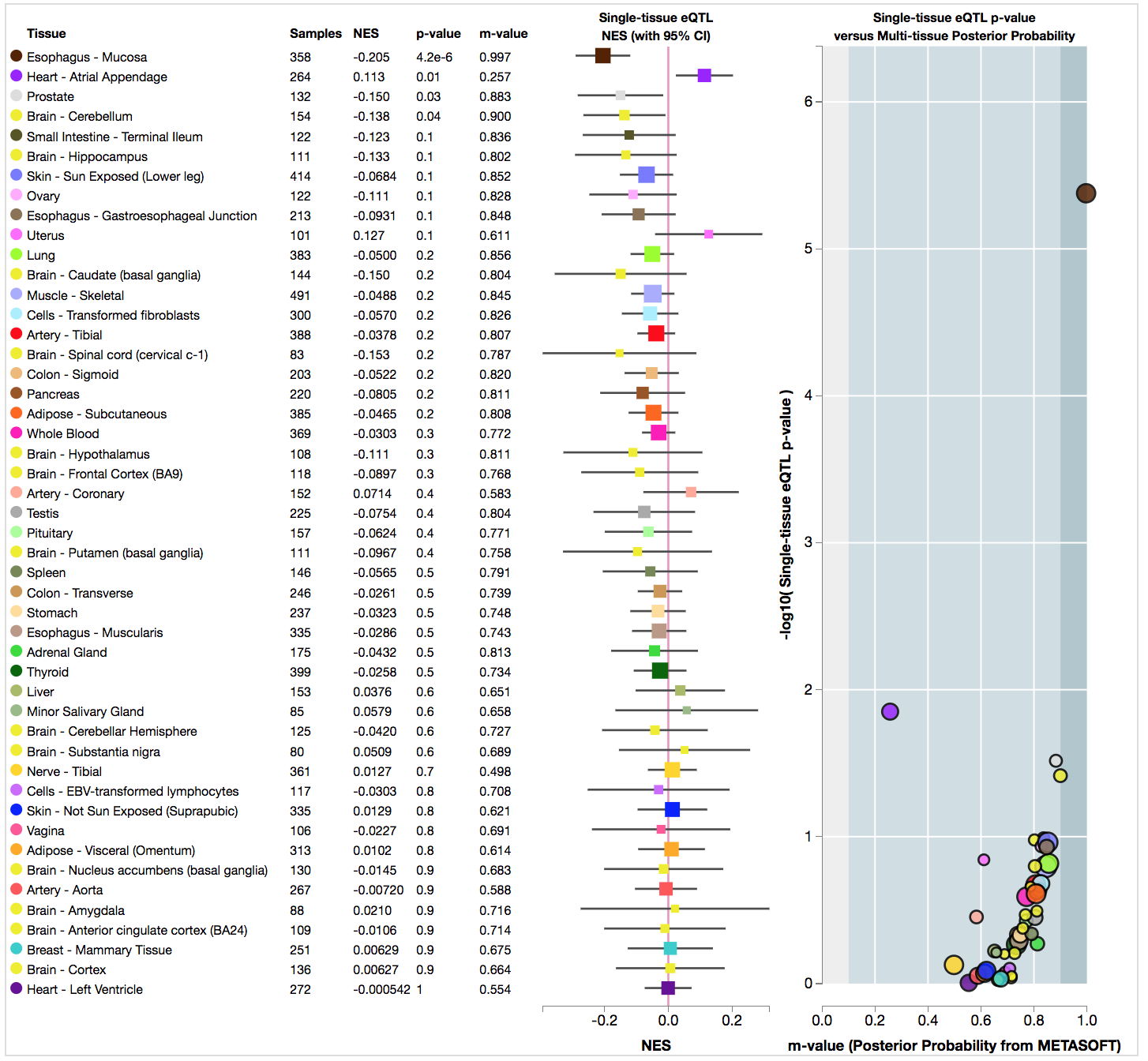

**Supplementary Figure 4:** rs35278 & rs165940 mapped with the haplotypes of *PDE4D* previously studied by Tomppo et al (17). Both the SNPs are in some degree of D′ LD with haplotype Block 14*, as determined by Hedrick’s multiallelic D’ (ref). a) Shows the whole LD haplotype block structure as determined by Tomppo et al (17). b) Close up of the specific region of interest, from our SNPs (within the red box) to Block 14 from Tomppo et al (17)(rs13190249, rs1120303, rs921942, rs10805515 and rs10514862).

a)

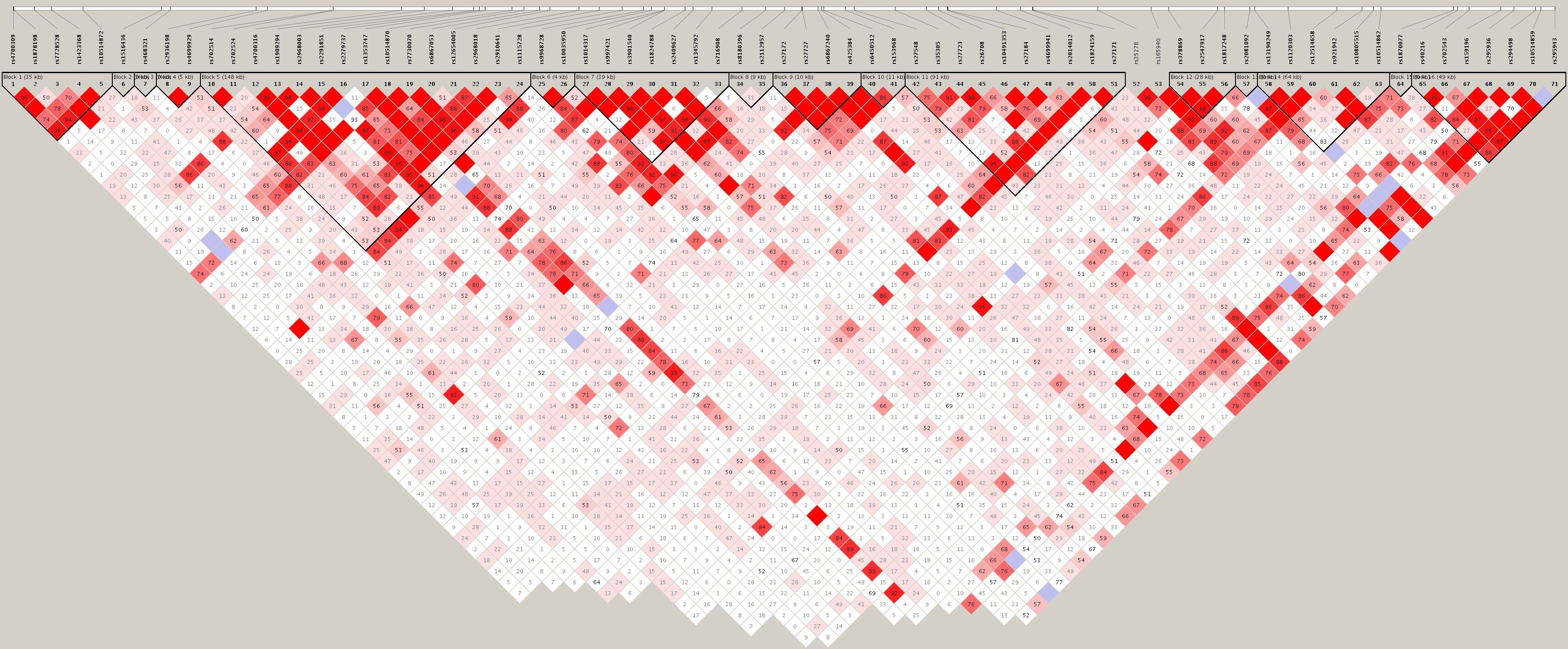

b)

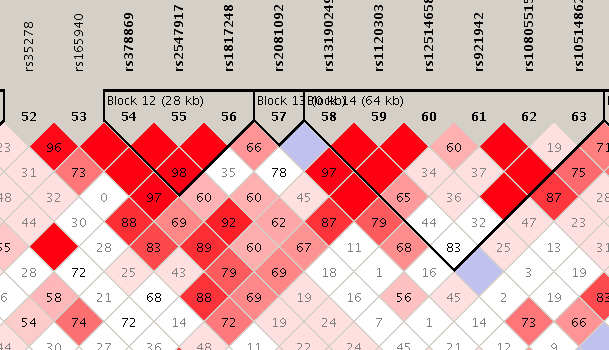

Both figures were generated through the program Haploview (18).

* Haplotype Block 14 here corresponds to Haplotype Block 15 reported by Tomppo et al (17). However, SNPs from another block were not successfully genotyped in that study, hence the numerical difference between block numbers in these two studies.
